# Gut-on-Chip Methodology Based on 3D-Printed Molds: A Cost-Effective and Accessible Approach

**DOI:** 10.1101/2025.01.29.632980

**Authors:** Elise Delannoy, Aurélie Burette, Sébastien Janel, Catherine Daniel, Alexandre Grassart

## Abstract

Gut-on-chips (GoC) represent a disruptive technology with great potential to understand the underlying mechanisms of gut health and pathology. Researchers who want to implement this approach, can either use expensive commercial microfluidic chips or build them from scratch in their lab. However, the design of such devices demands specific technical skills and expertise in computer assisted design (CAD). Additionally, the fabrication of the master molds for the chip production is very costly, time consuming and requires dedicated microfabrication facilities. Thus, the diffusion of these models in biology and health research laboratories remains limited due to this technological complexity and lack of affordability. In order to break these two bottlenecks, we present here the 3DP-µGut with open access designs and a simple fabrication approach based on a commercial 3D printer intended for general users. To ensure a good reproducibility with a sufficient number of available replicates for biological experiments, the method has been optimized to allow the production of multiple GoC per batch. The chips were also improved for confocal live imaging microscopy analyses. The chips are designed to be compatible with different types of microfluidics pumps, from stand-alone to completely integrated instrumentation. As a proof of concept, Caco-2 cells were seeded inside the fabricated GoC to validate their biocompatibility and functionality. After 7 days of maturation, cells self-differentiated in a 3D epithelium resembling *in vivo* expected structures. In summary we proposed a low-cost and open access GoC design and fabrication method with medium throughput. These results demonstrate the advantage of using 3D SLA printing to accelerate GoC implementation for gut physiopathology research.

## 3 INTRO

Organ-on-Chips (OoC) are rapidly emerging as new standard *in vitro* models. Their ability to mimic human organs in terms of structure, functionality and mechanical or chemical stimulation makes them a powerful and versatile research tool. Their implementation requires interdisciplinary skills combining engineering, materials science and biology approaches to reproduce living and functional tissues. OoC technology is particularly relevant for studying intestinal function and pathology due to the complex mechanical and 3D cellular structuration of the gut. In recent years, Gut-on-Chip (GoC) models outperformed conventional *in vitro* culture systems in mimicking the gut *3D* microstructure and its specific dynamic microenvironment [1]. Nonetheless, widespread adoption of these models in biological laboratories has been delayed by the technological challenges associated with their fabrication.

The historical and conventional method for constructing OoC devices, including GoC models, consists in the soft lithography of polydimethylsiloxane (PDMS) on SU-8 and silicon-based molds. PDMS is a widely used and well validated material for tissue engineering and OoC [2], it can be easily implemented in any lab for rapid microfluidic chip production. On the other hand, the fabrication of dry etched silicon or SU-8 mold requires heavy equipment and extensive specialized training. This process is very precise, achieving a nanometer resolution, but is yet very expensive and time consuming due to protocols with numerous steps [3]. Another critical limitation is its lack of geometrical liberty for the structures of such molds: they are restricted in height and only allow planar patterns with a rectangular cross section. Several new approaches have been proposed to simplify mold production and make it more accessible for biologist researchers. Subtractive methods, like micro milling of plastics and aluminum for instance [4], [5], [6], or laser cutting [7], [8] are more affordable and show good potential for microfluidic applications but are limited by their relatively low resolution (0.1 mm) and subsequent surface roughness.

In the past decade, SLA 3D printers have gathered a strong interest from the microfluidic and organ-on chip community because of their inexpensiveness and high resolution [9], [10], [11]. They rely on an additive manufacturing method, the photopolymerization of a photocurable ink in successive layers, allowing the creation of intricate microstructures. If 3D printed molds are already used for many microfluidic applications [12], [13], they still face some challenges for OoC approaches. Several groups have proposed 3D printed mold to build OoC devices [14], [15], [16], [17] but some encountered resin toxicity issues as uncured monomer can leachate inside the PDMS during molding [18].

We propose here a successful method, the 3DP-µGut, to build a GoC device at medium throughput based on 3D printed molds with general user equipment: a commercial 3D printer and resin. The chip layout was inspired from the largely adopted and successful design of Ingber and colleagues [19] divided in two connected parts with a top and bottom chamber. The channel was modified in a curved conduit to better accommodate the gut spatial geometry. The molds are designed for a good reproducibility and are reusable multiple times without alteration of the printed patterns. Each pair of molds (for the top and the bottom parts) allows the fabrication of nine GoC per batch. The final devices are optimized for imaging purposes thanks to a thin PDMS layer on the bottom side. Additionally, the microfluidic chip has been simplified and optimized to be compatible with plug and play solutions, implementable with minimal training, and is also suitable for more open instruments in the case of specific applications. 3DP-µGut devices were seeded with Caco-2 cells, a widely used intestine epithelial model, to assess their biocompatibility and their capacity to mimic the gut 3D architecture as observed in state-of-the-art models [20]. This fabrication process is simple, robust and affordable in producing Gut-on-chip devices for gut health studies.

## 4 MATERIAL AND METHODS

### 4.1 COMSOL simulation

Shear stress inside the 3DP-µGut was evaluated using COMSOL (COMSOL Multiphysics 5.5, COMSOL Inc.). The simulated curved channel was 11mm long, 750 µm wide and 750 µm high with a spatial period of 3 mm. The steady state was calculated with laminar flow physics. Navier stokes equations for incompressible flow were applied. The fluid flow was set at 60 µL/h and a non-slip condition was applied to the channel walls.

### 4.2 Mold printing, post-treatment, characterization

3DP-µGut molds were designed using Fusion 360 (Autodesk). For printing, STL files were uploaded in Preform software (Formlabs, USA) using “adaptable” parameter and prints were directly performed on a build platform 2 using a Form3 SLA 3D printer (Formlabs, USA). After careful removing from the build platform 2, prints were washed to remove excess uncured resin with IsoPropyl Alcohol (IPA) for 20 min using FormWash (Formlabs, USA) and dried with air, followed by 60 min of UV curing at 60°C using FormCure (Formlabs, USA). The printed mold was then salinized by nebulization of Trichloro(1H, 1H, 2H, 2H-perfluorooctyl) Silane (Sigma Aldrich, USA) under a vacuum during 2 hours. and heated at 90°C for 1h. The molds were imaged and measured with VHX-X1 microscope (Keyence, Japan). The height of the printed pattern was measured by focusing successively on the bottom and the top of the pattern. The 3D reconstruction of the ROI and measures of the height profile was calculated using the 3D module (Keyence, Japan).

### 4.3 Chips fabrication and assembly

For chip fabrication, a ratio of 1:10 w/w of the curing agent and PDMS pre-polymer (Sylgard 184, Dow Corning, USA) was used. For the bottom molds, plexiglass sheet and a 500 g weight were placed on top to control the height of the resulting PDMS slab. PDMS was cured overnight at 65°C. The PDMS was then carefully peeled of the molds surface, cut in individual parts.

The top compartments were exposed to O_2_ plasma at 100 W, 50 kHz (Cute, Femto Science, South Korea) and bonded with a Polyester (PET) porous membrane in the middle (ipCELLCULTURE™ Track-Etched Membranes, pore size: 8 µm, it4ip, Belgium) previously treated by 0_2_ plasma 100W, 50kHz and 5% APTES (Sigma Aldrich, USA) at 75°C for 20 min. The bottom compartments were then exposed to O_2_ plasma 100 W, 50 kHz and aligned with the tops to complete the gut chip. The chips were washed successively with 70% ethanol and milli Q water and then air dried. Eventually, chips were exposed to UVO (UVO cleaner, Jelight Company inc., USA) during 20 min for sterilization and stored in Petri dish before use.

### 4.4 Atomic Force Microscopy

Experiments were performed on a Bruker BioScope Resolve AFM coupled to a Zeiss Observer.Z1 optical microscope. All PDMS surfaces were scanned in air in contact mode using Bruker SNL-C probes at 0.5 Hz in NanoScope 9.4 software. Average (Ra) and RMS (Rq) roughness was analyzed using the corresponding data analysis software for 100 x 100 µm scans.

### 4.5 Cell Culture

Caco-2 cells (clone TC-7) were obtained from Sansonetti lab. Cells were grown in Dulbecco’s Modified Eagle Medium (DMEM, Gibco) supplemented with 20% FBS (fetal bovine serum, Fisher Scientific, France) 100 U/mL penicillin, 100mg/mL streptomycin and non-essential amino acid (Gibco) in 5% CO_2_ at 37°C.

### 4.6 Human Gut-on-Chip seeding and culture

The 3DP-µGut PDMS chips were coated overnight at 37°C, 5% CO_2_ with type I collagen (ThermoFisher, USA) and Matrigel (Avantor, USA) diluted in Caco-2 culture medium at respectively 30 µg/mL and 100 µg/mL. The channels were rinsed with cell culture medium and cells seeded at 2.10^6 cells/mL in the upper channel for 4 hours at 37°C, 5% CO_2_. After cell attachment, the top channel was gently washed using warm cell culture media. Chips were maintained statically overnight. For the chips under flow conditions, the top and/or the bottom were connected either to a microfluidic circuit actuated by Flow EZ pressure controllers (Fluigent, France), or an integrated and connected device OMI (Fluigent, France). They were cultured under 37°C and 5% CO2 conditions for 7 days as their static counterparts. The flow was set to 60 µL/hour. Cell culture media was renewed every 48 hours.

### 4.7 Immunofluorescence

3DP-µGut were fixed in PBS, 4% paraformaldehyde for 30 min at room temperature, permeabilized in PBS, 0.5% BSA, 0.2% Triton-X100 (Sigma-Aldrich) for 15 min at room temperature and blocked in PBS, 5% BSA (Sigma-Aldrich) and 2% FBS (Fisher Scientific, France) for 1 h at room temperature. GoC were then incubated with the primary antibody in PBS, 0.5% BSA, 0.2% Triton-X100 (Sigma-Aldrich) for 2 h at room temperature, rinsed 2 times for 5 min with PBS and incubated for 1 h at room temperature with the secondary antibody in PBS, 0.5% BSA, 0.2% Triton-X100 and rinsed as above. GoC were incubated in PBS, 1 µg/mL DAPI (Thermofisher, USA) for 20 min at room temperature, rinsed twice for 5 min with PBS and stored at 4°C until observation. The antibody primary and secondary antibody were:

| Antigen | Dilution | Catalogue # | Isotype | Manufacturer |
| --- | --- | --- | --- | --- |
| **Primary antibody** |  |  |  |  |
| CD324(E-Cadherin) | 1/50 | 610182 | Mousse | BD Biosciences |
| Villin 1 | 1/100 | 13463197 | Rabbit | Thermofisher |
| ZO-1 | 1/100 | 61-7300 | Rabbit | Thermofisher |
| **Secondary antibody** |  |  |  |  |
| Alexa 488 | 1/1000 | A32790 | Donkey anti Rabbit | Thermofisher |
| Alexa 568 | 1/1000 | A10042 | Donkey anti Rabbit | Thermofisher |
| Alexa 568 | 1/1000 | A11004 | Goat anti Mouse | Thermofisher |
| **Other** |  |  |  |  |
| Phalloidin Alexa 488 | 1/400 | A12379 |  | Thermofisher |
| Phalloidin Alexa 568 | 1/400 | A22283 |  | Thermofisher |
| Phalloidin Alexa 647 | 1/400 | A22287 |  | Thermofisher |

For transversal sections, 3DP-µGut chips were embedded in thick 5% agarose gel and cut with Compresstome® VF-310-0Z (Precisionary Instruments) in 400 µm thin segments. The slices were preserved and mounted with PBS on glass coverslip prior to imaging.

### 4.8 Microscopy

Vertical slices of the empty gut chip were imaged under a video-microscope Axio Observer Z1 equipped with a Prime95B camera (Zeiss). The maturation of the 3DP-µGut were followed by phase contrast imaging with an Eclipse TS2 inverted microscope (Nikon). Fluorescence images were taken using a SpinningDisk microscope equipped with a CSU-W1confocal scanner unit (Nikon) and objectives 10x CFI Plan Fluor (NA 0.30), 20x CFI Plan Apo Lambda (NA 0.75), 40x CFI Plan Apo (NA 0.95). Microscopy images were adjusted for brightness, color balance, and/or contrast uniformly across all pixels, as necessary, in image J (v1.46r, National Institutes of Health) [21]).

### 4.9 Permeability assay

After 7 days of culture, FITC-dextran 70 kDa was injected to the top channel, 10 µg/mL in cell culture media. Empty 3DP-µGut were used as a control. After overnight incubation at 37°C and 5% CO_2_, samples from the top and bottom were collected and stored in the dark at 4°C upon measurement. Fluorescence levels were measured in a VictorX3 Multilabel Plate Reader (PerkinElmer, USA) with excitation wavelength of 485-414 nm and an emission wavelength of 535-525 nm.

## 5 RESULTS

### 5.1 SLA 3D-Printing produces robust and reproductible molds for PDMS casting

The overall 3DP-µGut design consists of two channels separated by a porous membrane following a modified curved channels geometry replicating the intricacy of the human intestine and its physiodynamics [19], [22]. To optimize chip production, molds were created by pairs, one for the bottom channel of the chip (blue, Figure 1A) and one for the top channel (Red, Figure 1A). Each mold is composed of nine identical micropatterns shown in Fig1A, respectively 200 µm and 750 µm high. The design integrates pillars of 1.2 mm diameter for the inlets and outlets access to avoid fastidious manual punching and its associated variability. To confirm that the resulting shear stress is in physiological scale, the medium flow inside the device was numerically modeled (Figure 1B). The simulated shear stress is distributed across the channel, perpendicularly to the flow direction, ranging from 1 to 10.10^-4^ Pa (0.001 to 0.01 dyne/cm2).

**Figure 1:**
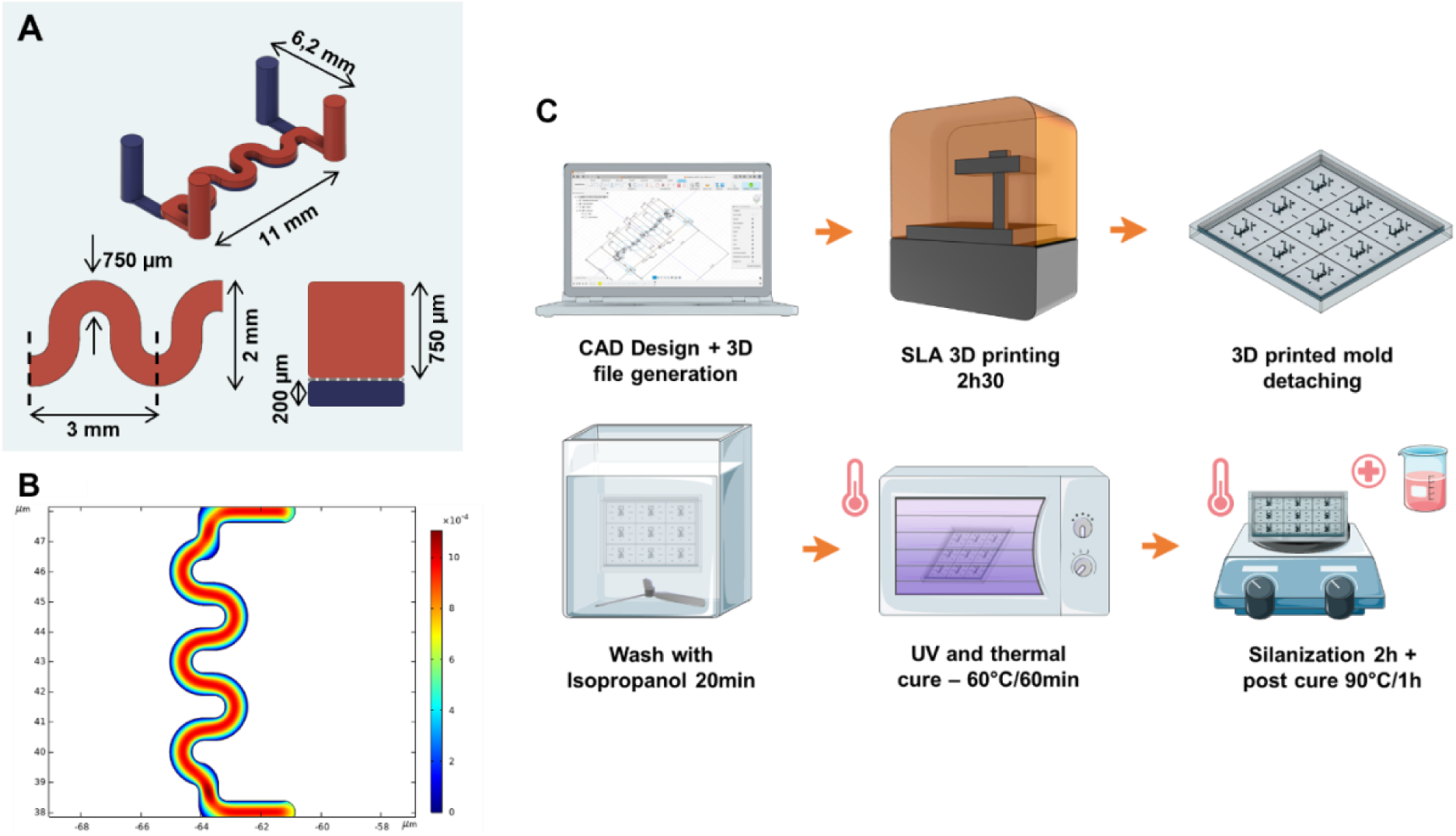
Design, Simulation and Fabrication process of the 3D Printed mold. A) Schematical view of the microfluidic channels. From top left to bottom right: 3D view, top view, vertical cross-section. The TOP channel is in red and the bottom channel in blue. B) Simulated shear stress inside the chip geometry, the results are expressed in Pa. C) Fabrication process for the 3D printed molds and post treatment steps.

The master molds were then fabricated using a commercial desktop SLA printer (Form3, Formlabs). The printed pieces were then post-treated according to the protocol described Figure 1C. Briefly, they were rinsed with isopropanol, post cured under UV and silanized to ensure quick demolding and preventing stickiness of the PDMS to the resin molds. In comparison to the conventional microfabrication techniques with soft lithography, this 3D printing-based fabrication protocol takes less than a day, can be easily implemented in a biology lab and is cost effective (Table SUPP1).

We validated the 3D printing technique as a robust alternative for mold fabrication by analyzing the reproducibility and resolution of the fabricated structures. Molds were characterized by measuring the height of the patterns for the bottom and the top mold with a high-resolution digital microscope (Figure 2C). Measurements show a low variability between the patterns from the same mold with meanSEM_Top_ = 4.4 µm and meanSEM_Bottom_ = 1.74 µm, and reproducibility between different molds with SD_Top_ = 15.3 µm and SD_Bottom_ = 7.4 µm from one mold to the other. The rugosity of the 3D printed mold was evaluated by AFM on PDMS slabs molded on either a SU-8 mold or a 3D printed mold (Figure 2D). As, expected, the rugosity of the 3D printed mold Ra=270nm is higher than is counterpart the silicon wafer with Ra = 14 nm (Table 1). However it remains low enough to guarantee optical transparency for cell observation by confocal microscopy, and our molds exhibit similar roughness to what was previously shown for similar mold printing techniques [23].

**Table 1:**
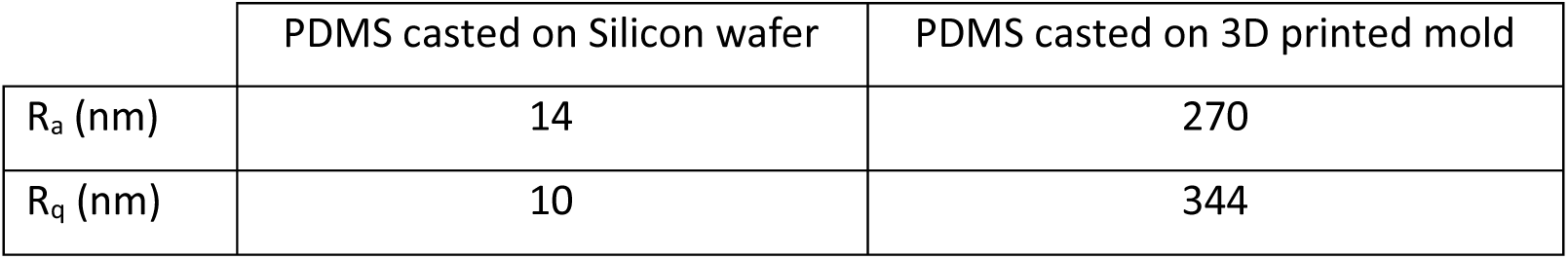
Roughness measurement for PDMS parts casted on Silicon wafer molds and 3D printed molds.

**Figure 2:**
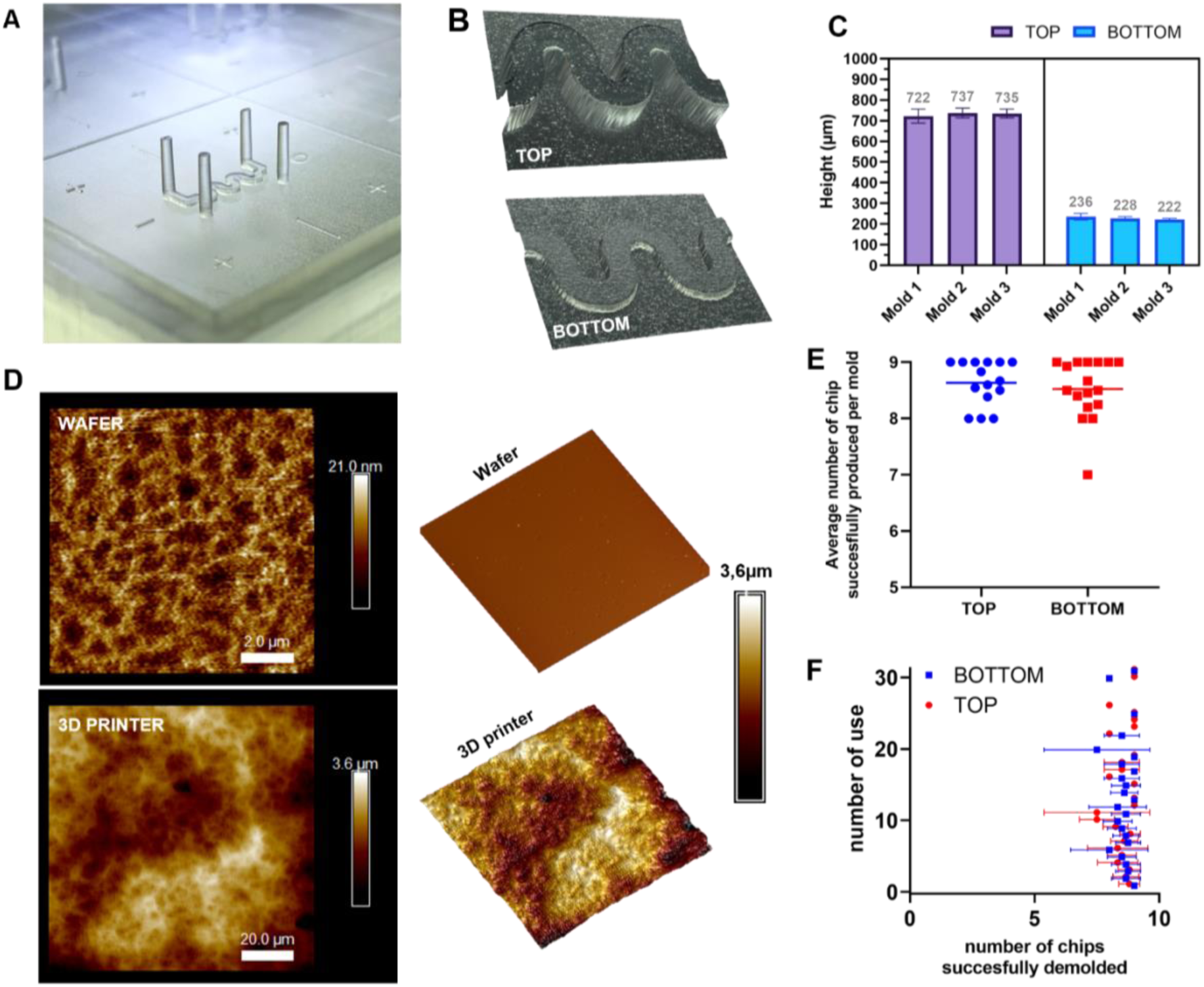
3D Printed mold characterization. A) 3D reconstitution of the printed patterns imaged with a surfaced microscope. B) close up photo of one of the printed patterns for a TOP mold. C) Channel height measurements within the same mold of between different batches. D) Topography of PDMS slabs casted on either a silicon wafer or a 3D printed mold. E) Graph of the average number of chips produced per molds, one point is one iteration. F) Evaluation of the performance of the molds across time.

Next, we assessed the durability of molds. To do so, we followed the performances of each mold individually for 3DPuGut fabrication. The results show an average successful demolding superior to eight individual chips for the wide majority of the produced molds (Figure 2E). Interestingly, the top and the bottom molds provide a continuous high rate of successful demolding even after thirty uses, proving the robustness of the 3D printing technique to fabricate microfluidic molds (Figure 2F). However the successive PDMS curing at 65°C appeared to have a deforming effect on the 3D printed pieces (Figure SUPP 1) with a light incurvation of the mold base. To prevent this deformation and improving mold longevity, weights were added on top of the molds during PDMS curing.

### 5.2 PDMS chip assembly and optimization for high resolution imaging

PDMS was cast onto the 3D printed pieces to fabricate the 3DP-µGut devices. After demolding, the PDMS slabs were cut in individual half gut chips. Top and bottom parts were then assembled together with a PET membrane in between thanks to surface treatment with APTES, plasma exposure and contact bonding (Figure 3A). The resulting 3DP-µGut is presented in Figure 3B and 3C. On the vertical cut of the chip (Figure 3D), we can observe a good curing and faithful reproduction of the design geometry with sharp angles. Some undulations are visible on the sides of the top channels due to the layer processing of the 3D printer and its resolution in Z (25 µm) but this does not constitute an issue as the cells will be cultivated on the porous membrane and not on the walls. PDMS was molded on a new or an old (>25 uses) mold to challenge the durability of the 3D printed structures. The results are shown in Figure SUPP2 with no visible alterations of the mold.

**Figure 3:**
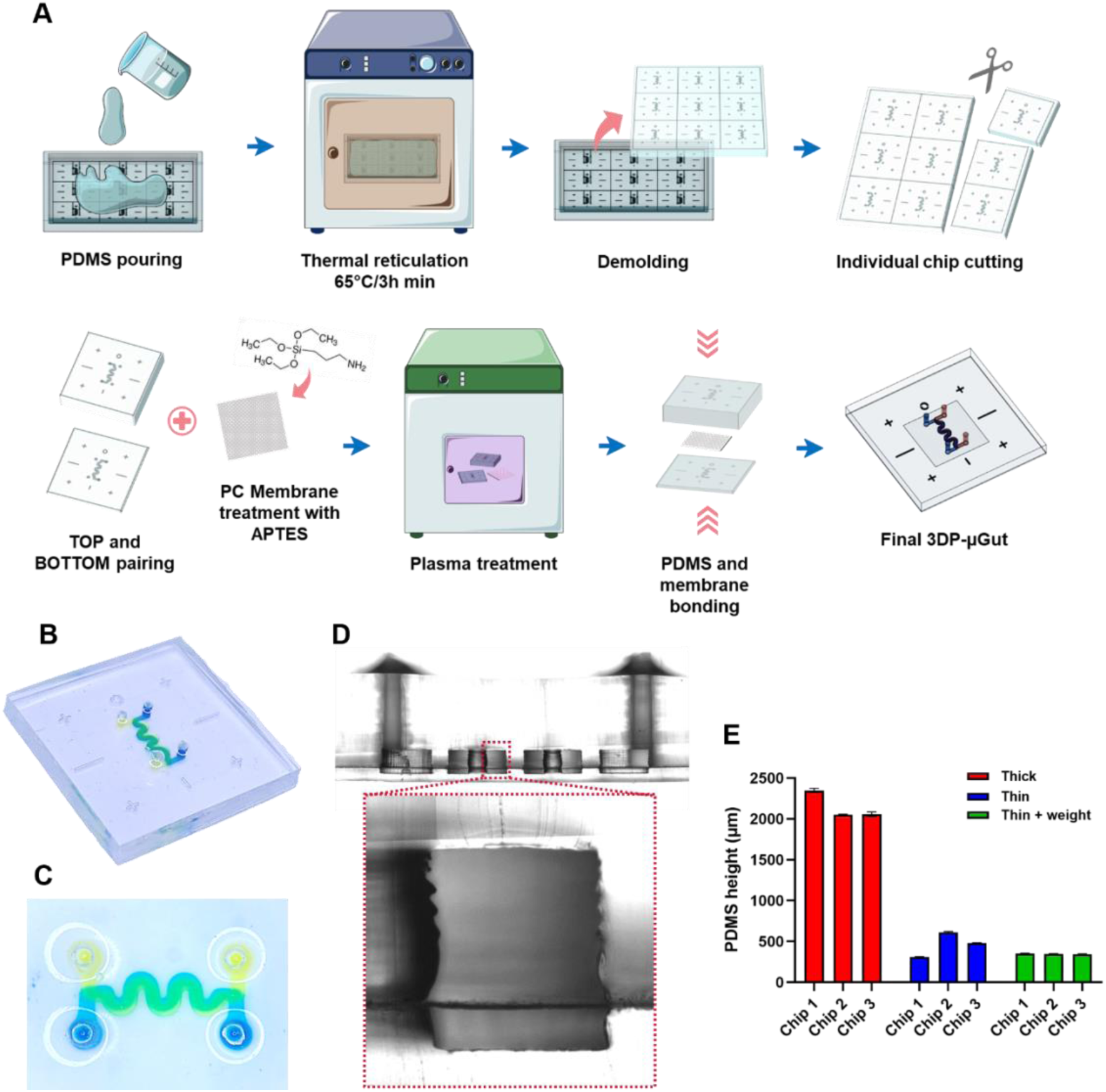
3DP-µGut assembly and optimization. A) Assembly steps for the 3DP-µGut. B) and C) views of the final 3DP-µGut device, top channel in yellow and bottom channel in blue. D) Cross section of the resulting PDMS chip. E) Thickness measurement of the PDMS layer under the BOTTOM channel depending on molding techniques.

Additionally, in order to optimize the devices for high-resolution imaging we needed the thinnest layer of PDMS as possible on the bottom side. To achieve submillimeter thickness, the liquid PDMS was mechanically pressed against the bottom mold with a PC flat square and thermally cured with a weight on top to ensure a controlled height. Figure 3E shows the differences in height measured depending on the molding technique, no press, pressed without weight and pressed with weight. The plastic lid combined with the weight appears to be the best method to achieve both very low thickness around 300 µm and repeatability.

### 5.3 Caco-2 differentiate in villi-like architectures inside the 3DP-µGut

To validate the biocompatibility of our 3DP-µGut devices with intestinal cells we coated and seeded Caco-2 cells inside the top channel. The devices were then connected to a flow circuit with microfluidic pumps and a continuous flow of medium at 60 µl/h was applied. We followed the maturation of the epithelium under phase contrast microscopy. Figure 4 shows the change in the structuration of cells and morphology under the effect of physiological flow. The Caco-2 differentiated from a cell monolayer after one day into a 3D invaginated micro architecture at day 7. As a control, cells were kept under static conditions and no structuration of the epithelium in 3D was observed at Day 7 (Figure SUPP 3).

**Figure 4:**
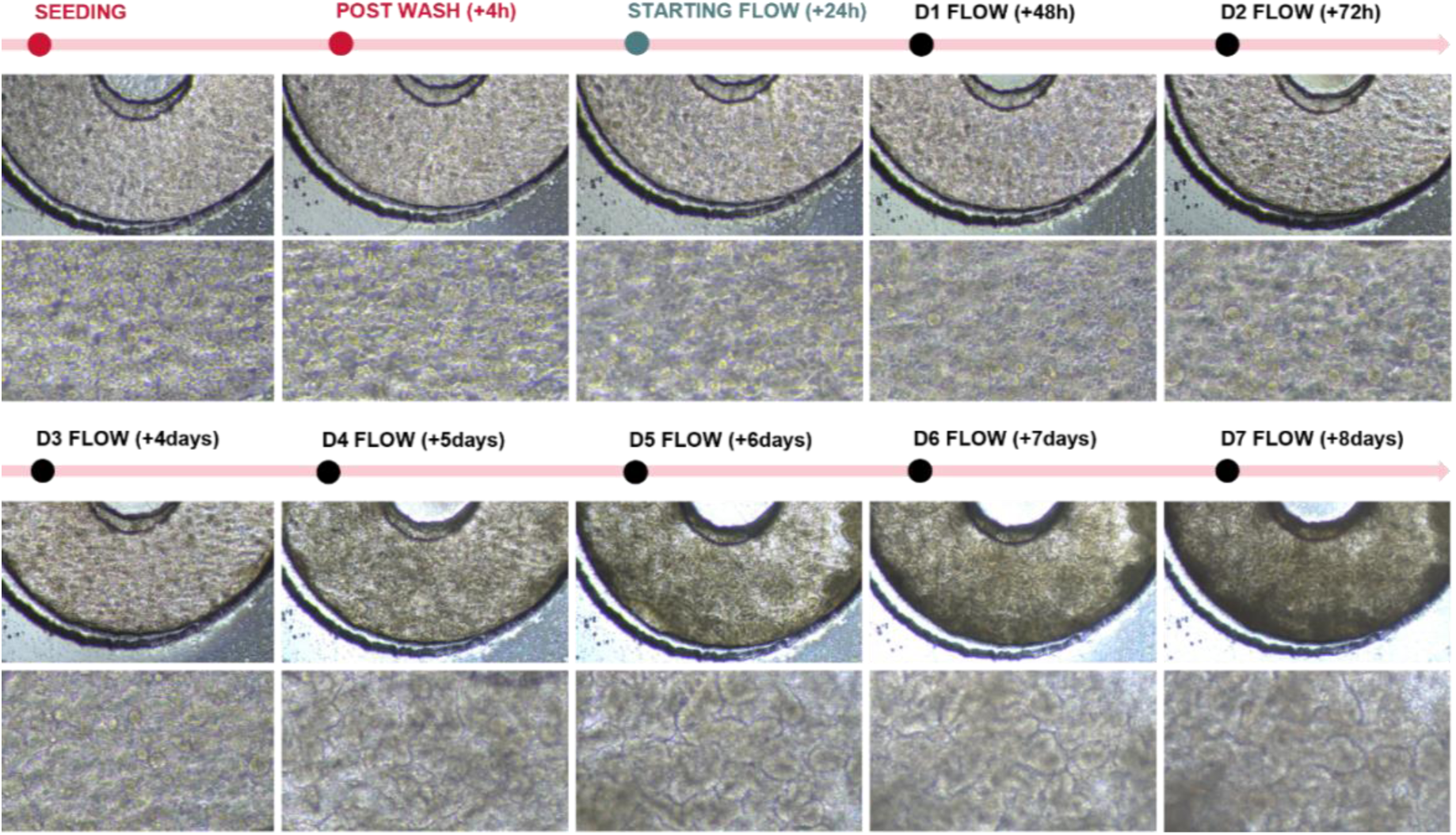
3DP-µGut maturation. Phase contrast imaging of the Caco-2 maturation inside the 3DP-µGut devices during one week under flow conditions, n=4.

To confirm further these structural observations, we embedded 3DP-µGut into agar, crosscut 400 μm sections and performed immunostaining. As observed previously, Caco-2 cells remain organized in a flat monolayer under static conditions, while villi and crypts-like structures are observed under continuous flow exposure (Figure 5A and 6B). Furthermore, the impact of the basal versus apical flow was investigated (Figure SUPP4). We observed that the basal flow is sufficient and necessary to achieve such cell organization, in contrary to the apical flow which is not sufficient by itself for a complete structuration, which is consistent with anterior studies [20].

**Figure 5:**
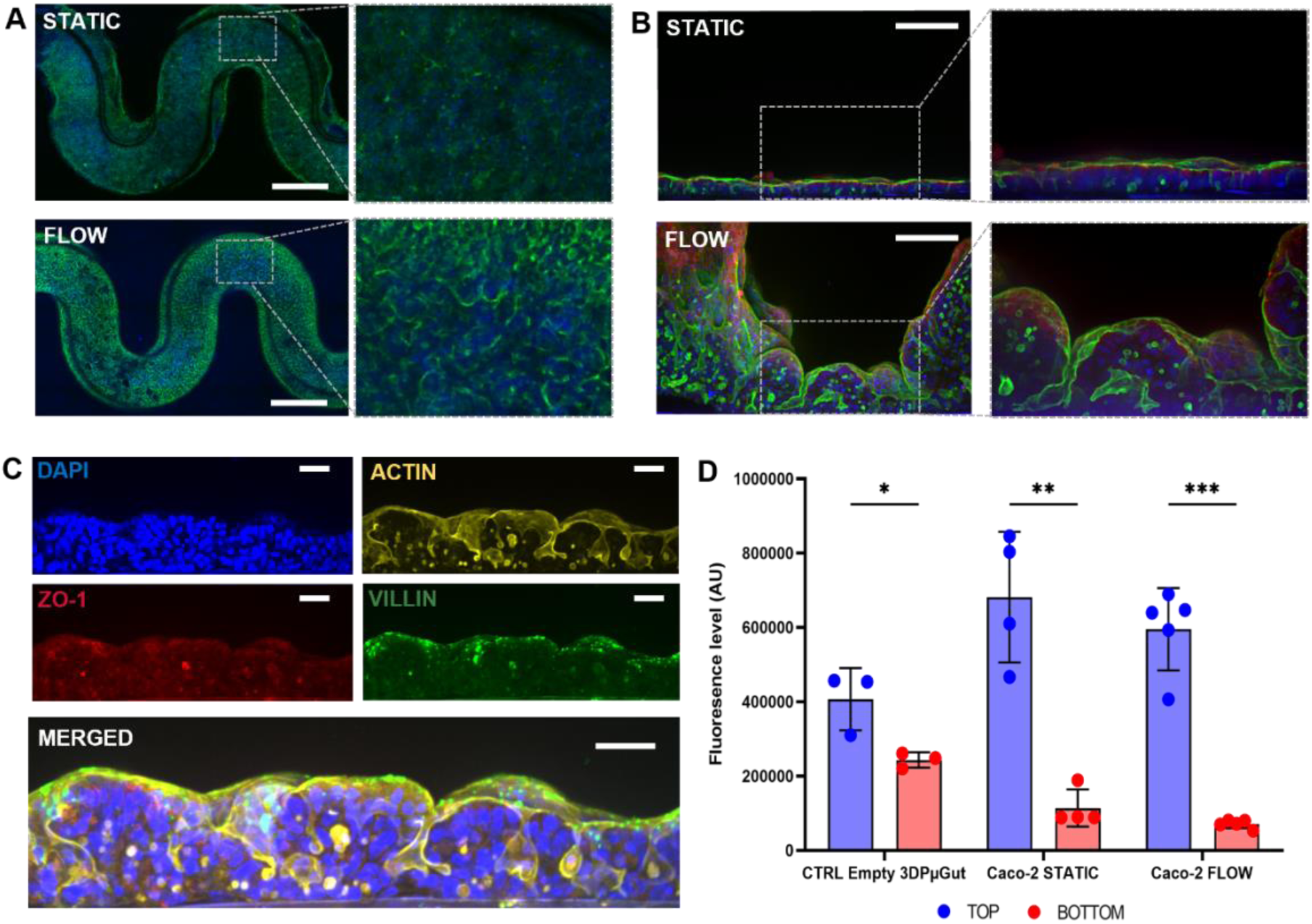
3DP-µGut 3D structuration. A), B) and C) Immunofluorescence staining of the Caco-2 cells inside the 3DP-µGut devices. n=3 A) Top view, Nucleus in blue (DAPI and Actin in green, bar=750µm B) Cross-section view, Nucleus in blue (DAPI), Tight junctions (Z0-1) in red and Actin in green, bar=100µm C) Cross-section view, Nucleus in blue (DAPI), Tight junctions (Z0-1) in red, Villin in green and Actin in yellow, bar=50µm. D) Permeability assay inside the 3DP-µGut devices in static or flow conditions, n≥3.

**Figure 6:**
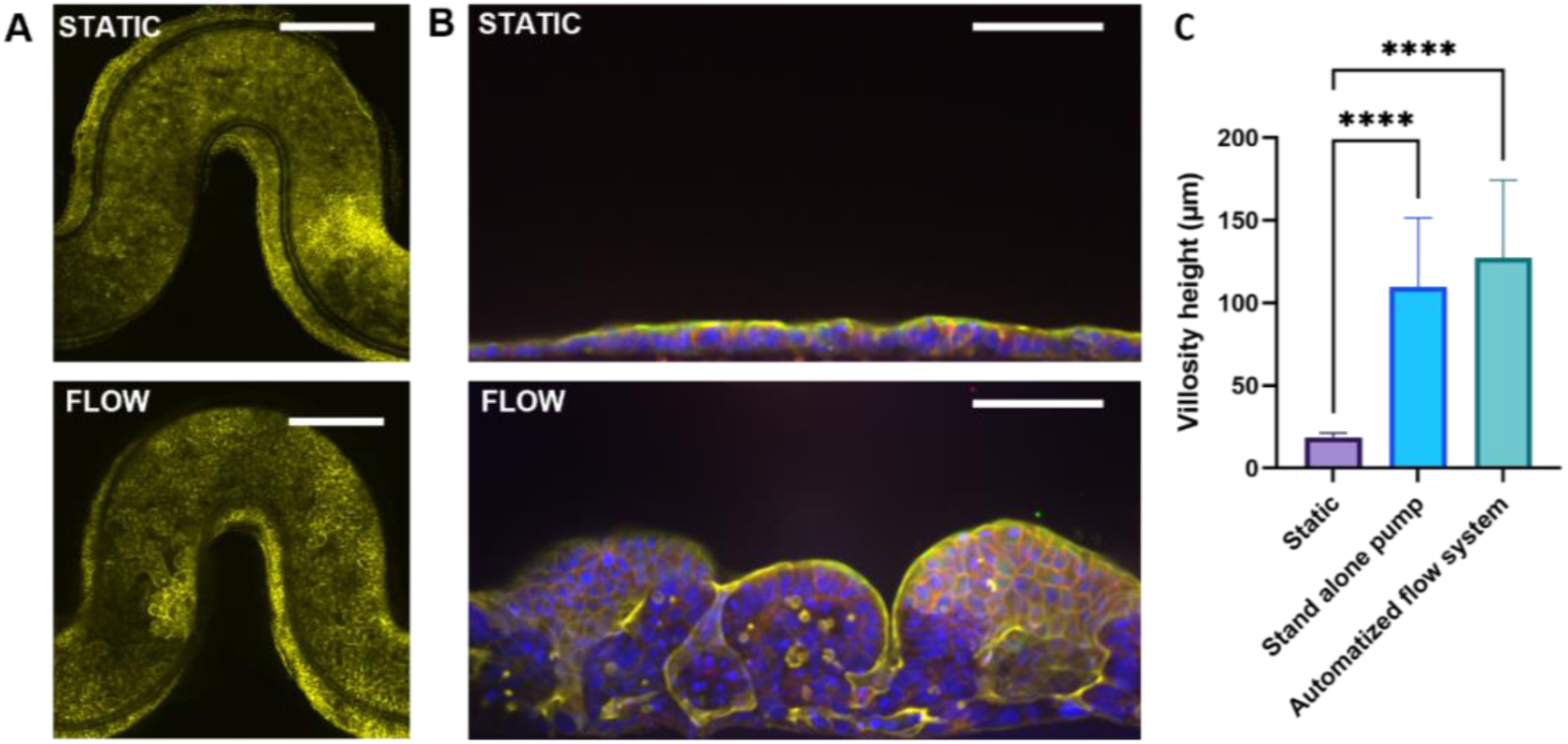
3DP-µGut Cell culture automatization with plug and play microfluidic platform. A) et B) Immunofluorescence staining of the Caco-2 cells inside the 3DP-µGut devices under static or flow conditions. n=2 A) Top view, Actin in yellow bar=750µm B) Cross section view, Nucleus in blue (DAPI), Tight junctions (E-Cadherin) in red and Actin in yellow, bar=100µm. C) Villi height measurement between static and flow conditions with stand-alone pumps or plug and play systems.

The functionality of the epithelium was evaluated with immunostaining, and cells exhibited tight junction markers ZO-1 as shown in Figure 5C. To assess the polarization and the differentiation of the Caco-2 cells the villin was also marked. As expected, the 3DP-µGut clearly expresses villin in higher concentration at the luminal side (Figure 5C). The epithelial permeability was observed by measuring transport of 70 kDa FITC-dextran across the cell barrier. The results show a strong decrease in permeability between empty control chip and colonized 3DP-µGuts indicating an effective barrier after seven days in culture in either static or flow conditions (Figure 5D).

### 5.4 Automatization of the 3DP-µGut cell culture with a plug and play microfluidic platform

One of the bottlenecks for a wider adoption of OoC is the complexity of the microfluidic instrumentation. Keeping the circuit sterile while performing live imaging can indeed be a struggle with conventional microfluidic pumps, requiring disconnecting the chips from the fluidic circuit. To prove the adaptability of the 3DP-µGut device, we automatized the culture of the Caco-2 with OMI platforms, which provide an all-in-one solution for chip perfusion (Figure SUPP 5).

The 3DP-µGuts were cultured under the automatized setup following the same protocol of 60 µL/h flow for seven days with medium recirculation. Fresh medium was introduced every 3 days. The automatized protocol and instrumentation resulted in the same self-organization of the Caco-2 cells in 3D microarchitecture (Figure 6A and 7B) exhibiting villi-like structure higher than 100 µm, similar to what was obtained with conventional pneumatic microfluidic pumps (Figure 6C). These results display the integration and automation potential for the 3DP-µGut and its future applications.

## 6 DISCUSSION

This study demonstrated the effectiveness of SLA 3D printing for the fabrication of molds used in the production of gut-on-chip devices: the 3DP-µGuts. The printed molds exhibit high resolution, sufficiently for reproducing cell culture microchannels. Unlike traditional photolithography, SLA 3D printing offers increased flexibility in mold design and customization, especially regarding achievable high aspect ratio [24]. They can be manufactured more quickly and at lower cost, while maintaining adequate accuracy for biological applications. The fabrication of the 3DP-µGuts does not require an extensive training or dedicated facilities, allowing its rapid implementation in non-specialized labs [25].

The versatility of this method allows the channel designs to be tailored for a wide range of applications and even for other organ-on-chip systems such as lungs [14], breast [18], or blood vessels [26], with similar size requirements. The structural details of the 3D-printed molds depend on the resolution of the selected printer. While the manufacturer specifies an approximate resolution of 100 µm in the XY plane, other studies have reported the successful fabrication of 25 µm features using the same general-purpose desktop SLA printer [27]. More resolutive 3D printing technologies already exist and achieve resolution of 2 µm [28] or even lower at the nanometer scale [29]. But they are costly, require extensive training and are limited by the total size capability of the final printed pieces. Fast advancement of 3D printing will certainly open new avenues in the field of organ on chip technologies.

Although PDMS casting inherently limits the throughput of microfluidic chip production, its integration with 3D-printed molds enhances manufacturing scalability. The optimized molds developed in this study allow the simultaneous production of nine 3DP-µGuts per batch. Furthermore, inlet and outlet access ports were integrated directly into the mold design, eliminating the need for time-consuming manual punching. The rapid fabrication of molds facilitates their multiplication and enables parallelized production of 3DP-µGuts, further improving efficiency. 3D printed molds are recently proven to be also compatible for higher throughput fabrication methods such as hot embossing [30].

Another approach consists in using the advantages of SLA manufacturing by printing the microfluidic devices directly with a biocompatible resin. Although, it remains challenging in terms of optical transparency, leakage management, cell adhesion and cytotoxicity [9]. A Biocompatible PEGDA has been used to print transparent bio-microfluidic devices [31]. The layer by layer printing could also pave the way to in situ 3D structuration of hydrogels inside the printed microfluidic channels [32] for instance to recreate intestinal structures, similar to the gut-on-chip proposed by Nikolaev and colleagues [33].

Caco-2 cell line was used here as a proof of concept of the biocompatibility of the fabricated chips and their ability in reproducing the physiological conditions for enterocytes maturation and structuration in 3D. As a perspective, more advanced cell sources such as human induced pluripotent stem cells (hiPSCs) or adult stem cells derived from patients could be used, as demonstrated by others using similar PDMS based gut-on chip models [34], [35]. In addition, the flexible and adjustable design of the 3D printed mold facilitates the testing of various intestinal microenvironment configurations. Coupled together, they could pave the way to patient-gut-on-chips and personalized medicine [36]. Additionally, the ability to fabricate customized molds could open the possibility of modeling intestinal architectures specific to certain pathologies, offering new avenues for disease modeling and therapeutic development.

## 7 CONCLUSION

Here we described the fabrication process of the 3DP-µGut devices with 3D printed molds. We assessed the reproducibility of the method and characterized the surface and the longevity of the produced molds. We evaluated the assembly of the final devices and tested their biocompatibility by cultivating Caco-2 cells inside the 3DP-µGuts. The cells exhibited spontaneous 3D structuring in villi-like morphology under flow conditions. They displayed typical enterocytes maturation markers and formed an effective epithelial barrier. Finally, we show the integrability of the 3DP-µGut within a plug and play fluidic system. Through this work we emphasize the potential for SLA 3D printing in being a promising technology for the fabrication of molds for intestinal-on-chip devices. Its advantages in terms of precision and design flexibility allow pushing the limits of *in vitro* models.

## 8 ACKNOWLEDGMENT

The authors would like to thank Fluigent for the lending of two OMI platforms. We also thank Sophie Salomé-Desnoulez from BICeL - US 41 - UAR 2014 – PLBS for her guidance on microscopy protocols. We also thank Jerome Vicogne from the Center of Infection and Immunity of Lille for granting us the access to the Keyence microscope. We would also like to thank Fabrice Soncin from CNRS/IIS/Centre Oscar Lambret/Lille University SMMiL-E Project for the access to the COMSOL software. We also thank members of MoHMI team for comments and discussion on the manuscript. This research is supported by Atip-Avenir funding, MEL “Accueil de talents” funding and region Hauts de France “Start Airr”. The French government under the France-2030 program, the University of Lille and the Lille European Metropolis (MEL) are thanked for their funding and support for the project R-CDP-24-007-MOSAIC granted to AG.

## 9 AUTHORS CONTRIBUTIONS

E.D. designed and performed most of the experiments and wrote the manuscript. A.B. fabricated the microfluidic chips and participated in the Caco-2 cells culture. S.J. performed AFM experiments. C.D copyedited the manuscript. A.G. led the work, participated in its funding, designed experiments, and wrote the manuscript with E.D.

## SUPPLEMENTARY DATA

**Tableau SUPP1:** Comparison between microfabrication and SLA 3D Printing techniques.

| Master Mold Frabication Methods | Microfabrication | SLA 3D printing |
| --- | --- | --- |
| Required Equipment Cost | Clean room facilities: Lithography equipment or DRIE >1Millions € | 3D printer 5000€ |
| Materials | Silicium wafer + SU8 resin | Clear resin |
| Resolution | 1nm | Z 25µm, XY 100µm |
| Fabrication time | 2days | 3h |

**Figure SUPP 1:**
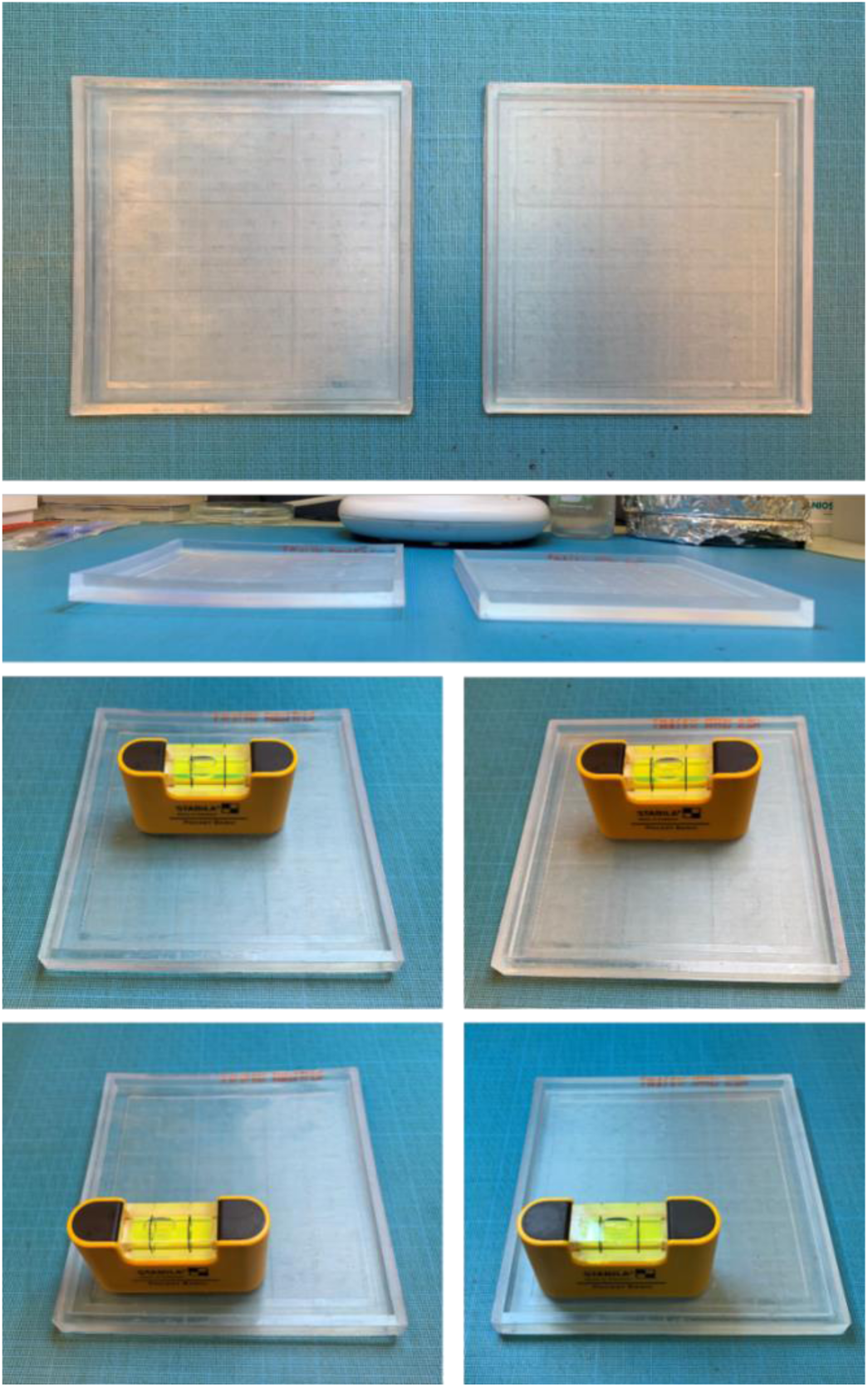
Comparison between a mold without (left) or with weight (right) on top after 30 heating cycles of 10h at 65°C

**Figure SUPP 2:**
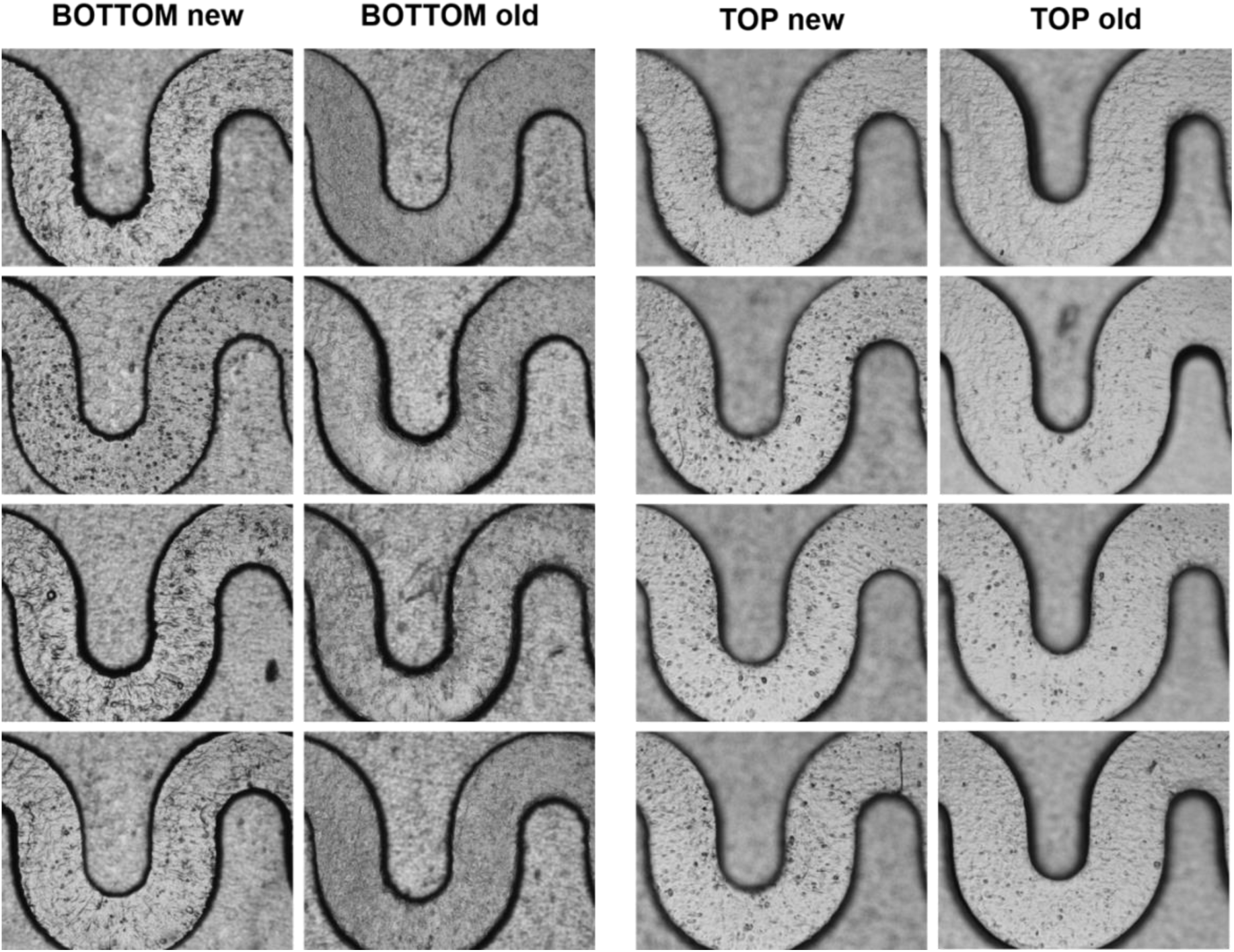
Imaging of PDMS channels casted on TOP or BOTTOM and new or old molds (25 uses)

**Figure SUPP 3:**
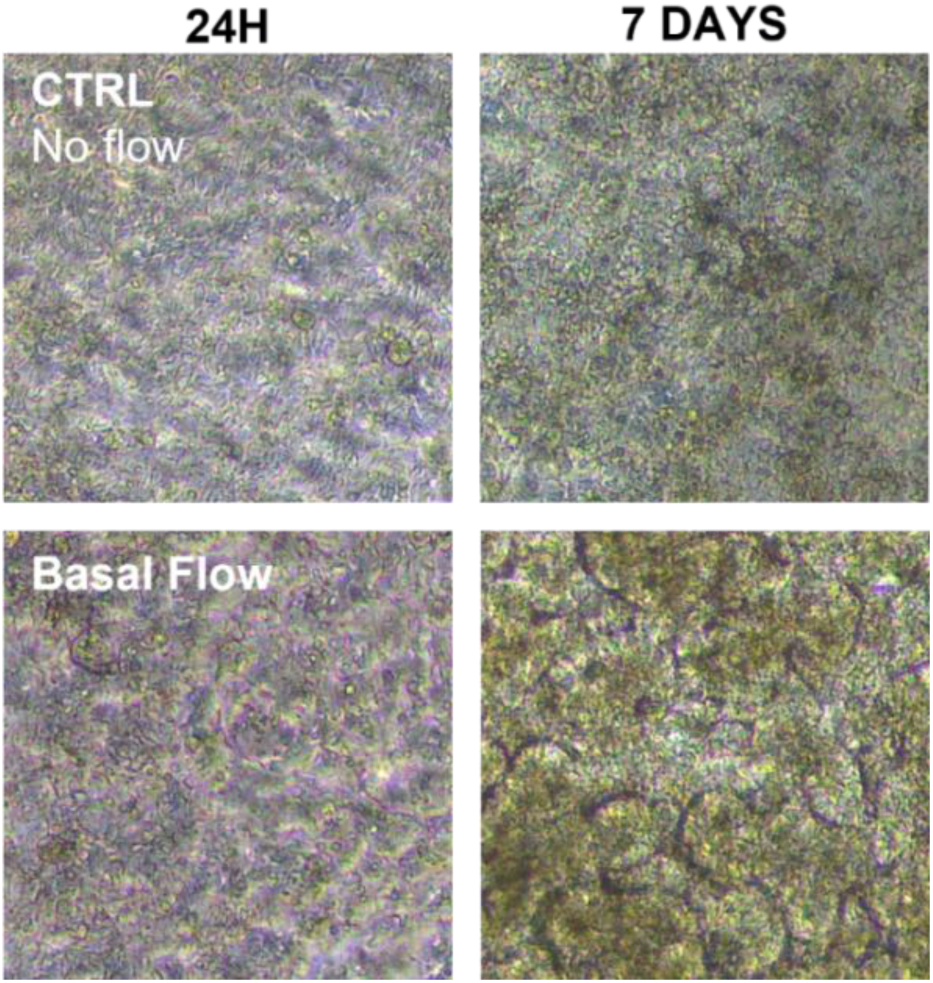
Phase contrast imaging of the Caco-2 maturation inside the 3DP-µGut devices during one week under static or flow conditions, n=3

**Figure SUPP 4:**
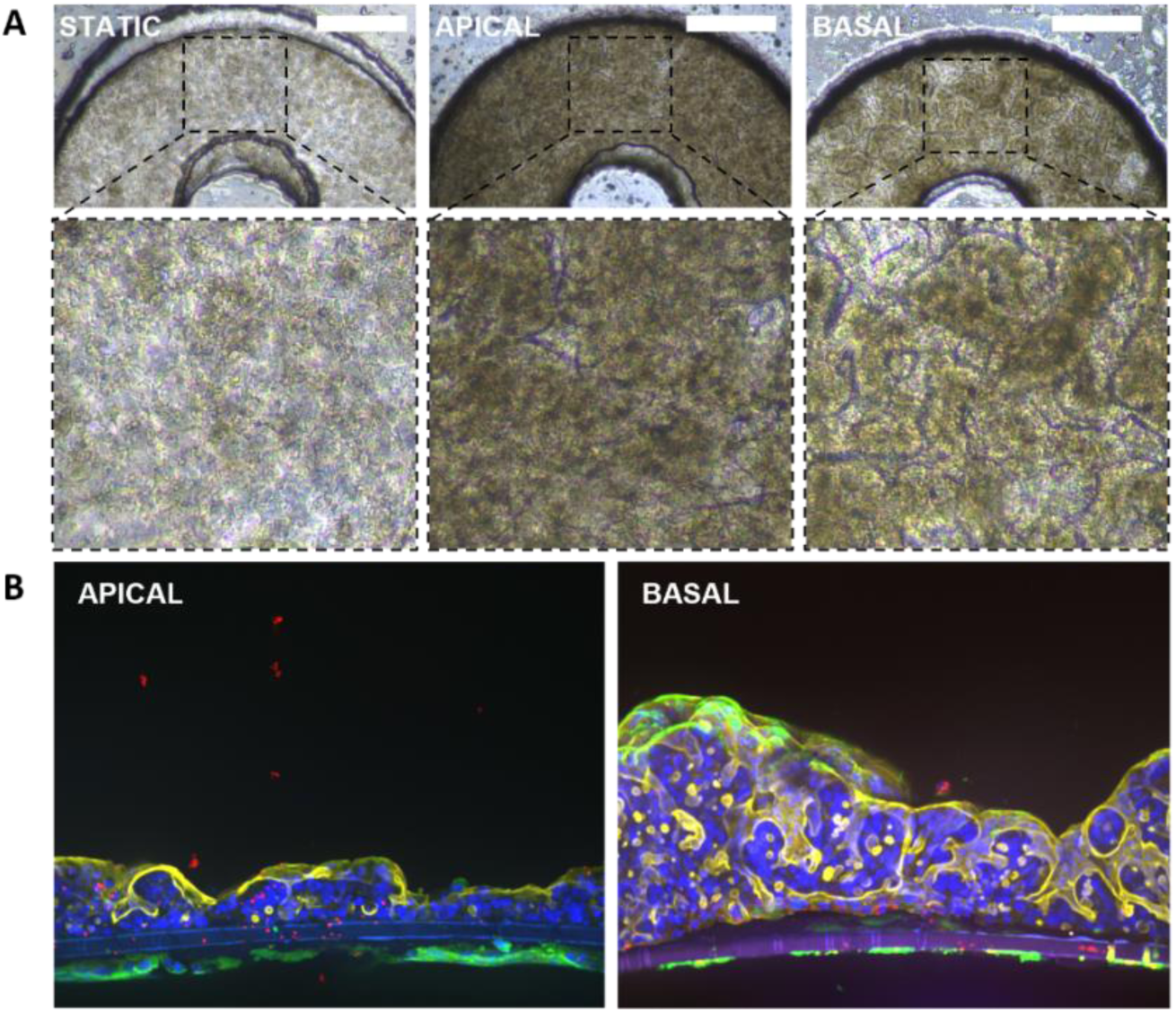
A) Phase contrast imaging of the Caco-2 maturation inside the 3DP-µGut devices during one week under static or flow conditions, n=3, bar=500µm.B) Immunofluorescence staining of the Caco-2 cells inside the 3DP-µGut devices. n=3 Cross-section view, Nucleus in blue (DAPI), cell junctions (E-cadh) in red, Villin in green and Actin in yellow.

**Figure SUPP 5:**
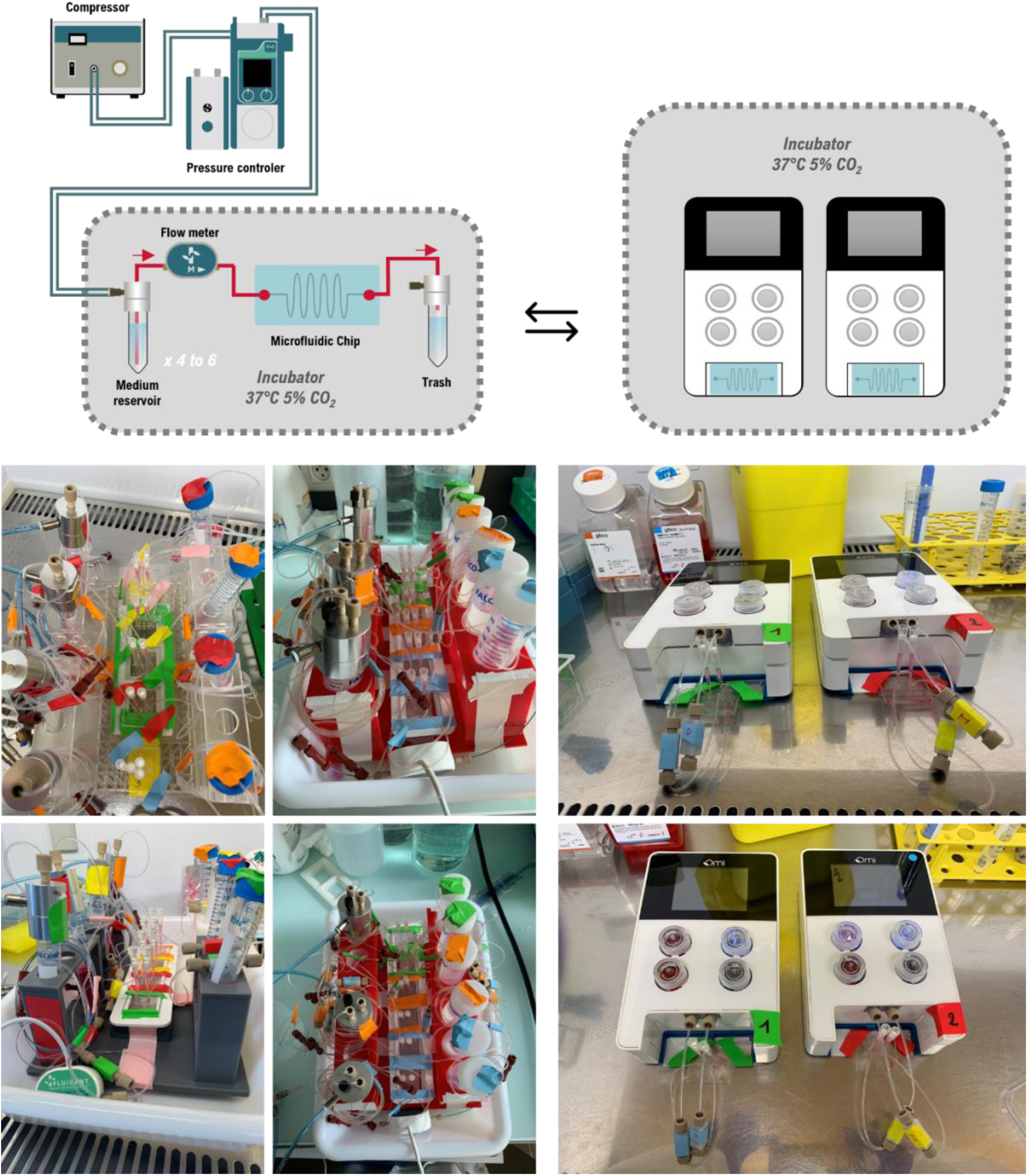
Fluidic circuit for the maturation of the 3DP-µGuts. On the left the circuit connected to the stand-alone pneumatic pumps Flow EZ, on the right the plug and play platform OMI

